# Autophagy in parvalbumin interneurons is required for inhibitory transmission and memory via regulation of synaptic proteostasis

**DOI:** 10.1101/2022.10.10.511533

**Authors:** Theodora Chalatsi, Laura M.J. Fernandez, Jules Scholler, Laura Batti, Angeliki Kolaxi, Leonardo Restivo, Anita Lüthi, Manuel Mameli, Vassiliki Nikoletopoulou

## Abstract

With the emerging role of the autophagic machinery in healthy brain development and aging, there is a pressing need to better characterize its functions in different neuronal populations, providing cellular insight into autophagy-related brain diseases. Here, we generated and characterized mice with conditional ablation of *atg5* in GABAergic neurons expressing parvalbumin (*PV-atg5KO*), mostly comprising fast-spiking interneurons, as well as Purkinje cells in the cerebellum. Using light-sheet microscopy to image PV neurons throughout the brain, we reveal that autophagy is required for the sustenance of Purkinje cells but not of PV-interneurons. Yet, proteomic analysis showed that autophagy deficiency in cortical and hippocampal PV-interneurons alters the proteostasis of key synaptic proteins, as well as the surface expression of glutamate receptor subunits. Consistently, hippocampal autophagy-deficient PV-interneurons exhibit reduced inhibitory neurotransmission and *PV-atg5KO* mice display excitation-inhibition imbalance in the hippocampus and memory deficits. Our findings demonstrate a neuronal type-specific vulnerability to autophagy deficiency, while also identifying PV-interneurons as cellular substrates where autophagy is required for memory.

## Introduction

In humans, genetic mutations resulting in diminished autophagic activity manifest as complex neurodevelopmental disorders, entailing epilepsy and impaired cognitive performance (Ebrahimi-Fakhari et al., 2016), or late-onset neurodegenerative diseases (Nixon, 2013). However, the roles of autophagy in the sustenance and synaptic transmission of diverse neurons remain insufficiently characterized.

In mammals, the developmental ablation of core autophagy genes, such as *atg5* (Hara et al., 2006; Kuijpers et al., 2021), a*tg7* (Komatsu et al., 2006) or *wdr45* (Zhao et al., 2015), leads to progressive and widespread neuronal death. In these studies, autophagy genes were ablated in multiple cell types of the brain, including progenitors and their derived neurons, astrocytes and oligodendrocytes, making it difficult to ascertain whether different types of post-mitotic neurons require autophagy in a cell-autonomous manner for their survival or sustenance.

Beyond neuronal survival, accumulating evidence suggests that autophagy also regulates several aspects of the physiology of glutamatergic neurons. These include the development and maturation of excitatory synapses, by facilitating the pruning of their dendritic spines (Tang et al., 2014), long-term synaptic depression (LTD), through the degradation of post-synaptic proteins (Compans et al., 2021; Kallergi et al., 2022; Pan et al., 2021) and basal excitatory neurotransmission, by regulating presynaptic ER-homeostasis and calcium stores (Kuijpers et al., 2021).

In agreement with its emergent synaptic functions, autophagy was also found to be essential in the hippocampus for memory and learning (Glatigny et al., 2019; Pandey et al., 2021). Yet the cellular substrates where autophagy is required for these complex behaviors remain elusive. This is largely due to lack of knowledge on whether autophagy also regulates the sustenance and synaptic properties of other neuronal populations, such as interneurons, which provide inhibition into the neuronal networks they are embedded in.

We hypothesized that the memory deficits associated with impaired brain autophagy may involve the dysfunction of Parvalbumin (PV)-expressing neurons. In the cortex and hippocampus, PV is mostly expressed by fast-spiking interneurons (Hippenmeyer et al., 2005). Despite comprising a small fraction of the brain’s neuronal network, these cells are characterized by unique morphological and molecular properties that make them key players of excitation-inhibition balance and GABAergic disinhibition between circuitries (Nahar et al., 2021). As well as being heavily implicated in neuro-psychiatric diseases (Kaar et al., 2019; Nakazawa et al., 2012; Ruden et al., 2021), PV-expressing neurons have high metabolic demands (Gotti et al., 2021; Inan et al., 2016; Kontou et al., 2021), rendering them potentially vulnerable to different types of stress (Page et al., 2019) including autophagy impairment.

To address this hypothesis, we generated and characterized mice with conditional ablation of a*tg5* in PV-expressing neurons. We demonstrate that Purkinje cells are selectively eliminated in the absence of functional autophagy, while the survival and sustenance of PV-interneurons are unaffected. Moreover, we reveal a pivotal role of autophagy in regulating the proteostasis and inhibitory neurotransmission of PV-interneurons, whose impairment has direct repercussions on the hippocampal network and on memory.

## Results

To determine the functional repertoire of autophagy in PV-expressing neurons, we crossed the *pv-cre* deleter mice (Hippenmeyer et al., 2007) with mice carrying floxed alleles of *atg5* (Hara et al., 2006). The resulting *pv-cre;atg5*^*f/f*^ progeny (referred to hereafter as *PV-atg5KO*) was generated at the expected mendelian ratios. Upon reaching adulthood (three-month-old),*PV-atg5KOs* presented a significant reduction in body weight when compared to their *atg5*^*f/f*^ littermates (referred to here as control) (Figure S1A, B). The two genotypes consumed similar amount of food, suggesting that differences in weight cannot be ascribed to altered feeding behavior in *PV-atg5KOs* (Figure S1C).

After crossing the *pv-cre* and *PV-atg5KO* animals to the robust Ai9 cre-dependent Td-Tomato (TdT) reporter strain (Madisen et al., 2010), we confirmed that a very high percentage of TdT-positive cells could be co-labelled using antibodies against PV in multiple brain areas (Figure S1D), as illustrated by representative images from the hippocampus and the somatosensory cortex (Figure S1E). Next, we FACS-sorted TdT-positive cells from *pv-cre;TdT*^*tg/+*^ (control) and *pv-cre;atg5*^*f/f*^*;TdT*^*tg/+*^ (*PV-atg5KO-TdT*) adult brains and performed JESS-Western blot analysis using antibodies against the autophagic substrate p62 (also known as Sqstm1). As shown in Figure S1F, p62 protein levels were significantly increased in neuronal lysates prepared from *PV-atg5KO-TdT* mice, when compared to the control, indicating a successful impairment of autophagic degradation within these cells. In line with this, we prepared protein lysates from the cerebellar cortex of adult (three-month-old) control and *PV-atg5KO* mice, a brain area containing a much higher proportion of PV-expressing cells. Western blot analyses indicated a significant accumulation of p62 (Figure S1G), as well as of another autophagy receptor, Alfy (also known as Wdfy3) (Figure S1H) in *PV-atg5KO* cerebellar lysates, both of which are known to accumulate when autophagic degradation is impaired. Moreover, analysis with an antibody against Atg5, recognizing the Atg5-Atg12 complex at 60KD, further confirmed a significant reduction of the Atg5 protein in *PV-atg5KO* when compared to control (Figure S1I).

In the cerebellum, PV is expressed by two distinct GABAergic populations: 1) the Purkinje cells, inhibitory projection neurons that co-express calbindin and are located in the Purkinje cell layer, and 2) the PV-interneurons found in the molecular layer. A previous study reported that autophagy is required for the maintenance of Purkinje cells, whereby the conditional ablation of *atg7* using a *pcp2-cre* deleter mouse led to their rapid degeneration (Komatsu et al., 2007). In line with these findings, immunolabeling cerebellar cryosections with antibodies against PV and Calbindin confirmed the drastic loss of more than half of the Purkinje cells in six-month-old *PV-atg5KO* brains (Figure 1A, B). Moreover, at three-months-of-age, the vast majority of Purkinje cells were immuno-reactive for active caspase-3 (Figure S1J,K), suggesting that the loss of these neurons is progressive. Consistent with the degeneration of Purkinje cells, three-month-old *PV-atg5KO* mice exhibited impaired motor coordination in a rotarod test, demonstrating a significant reduction in their mean latency to fall from the rotarod (Figure S1L) and across three consecutive trials (Figure S1M) when compared to control littermates. These data provide further support to the hypothesis that autophagy is required in Purkinje cells for their sustenance and for the expression of cerebellar-dependent behaviors.

**Figure 1:**
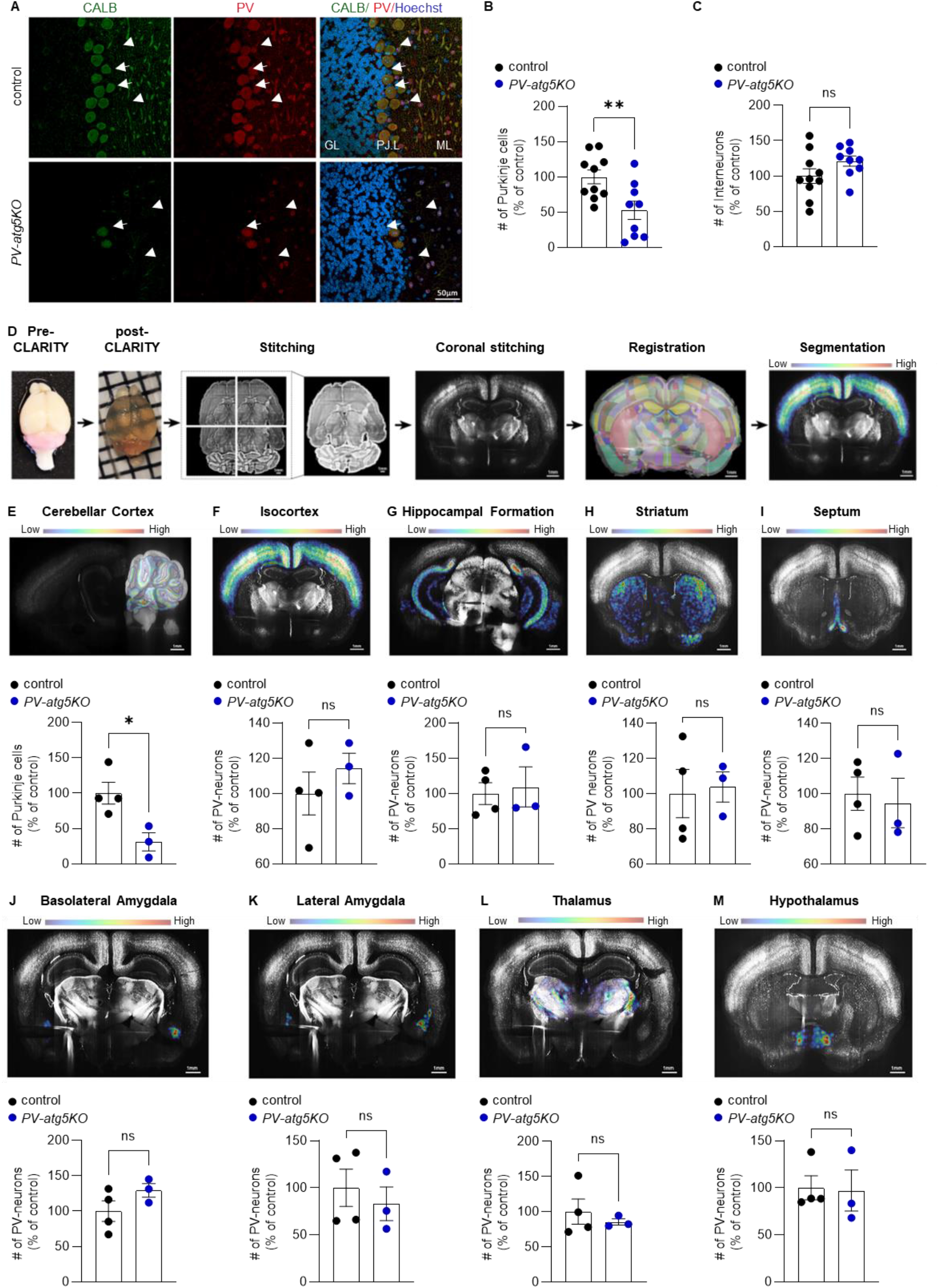
Selective elimination of Purkinje cells and not of interneurons upon genetic autophagy impairment. A) Representative confocal images of cerebellar sections from six-month-old control (*atg5*^*f/f*^) and *PV-atg5KO (pv-cre*^*+/-*^*;atg5*^*f/f*^) littermates. Immunostaining with antibodies against Calbindin (CALB, green) and Parvalbumin (PV, red), as well as with the nuclear dye Hoechst (blue) was performed. (Arrows: Purkinje cells, arrowheads: cerebellar interneurons, GL: Granular layer, PJ: Purkinje cell layer, ML: molecular layer). Scale bar: 50μm. B-C) Graphs showing the relative percentage of the number of PV-PJ cells and PV-interneurons, respectively, in the of cerebellar cortex of control (*atg5*^*f/f*^) and mutant *PV-atg5KO (pv-cre;atg5*^*f/f*^). Statistical analyses were performed on the number of cerebral sections that were counted using unpaired, two-tailed Student’s t test (**p=0.0091 t=2.941, df=17 and ^ns^p=0.1168 t=1.653, df=17 respectively). Bars represent mean values ±SEM. N=3 animals per genotype. D) Overview of the experimental pipeline for whole brain imaging and analysis. Three-month old control (*pv-cre;TdT*^*tg/+*^) and mutant *PV-atg5KO-TdT (pv-cre;atg5*^*f/f*^*;TdT*^*tg/+*^) littermates were analyzed. Scale bar: 1mm. E-M) Top rows: Representative heatmaps from the brain segmented masks from cerebellar cortex (E), isocortex (F), hippocampal formation (G), striatum (H), medial and lateral septum (I), basolateral amygdala (J), lateral amygdala (K), thalamus (L) and hypothalamus (M). Scale bar: 1mm. Bottom rows: Graphs showing the number of TdT^tg/+^ counted cells in the relative specific area, expressed as percentage of the average of the control. Statistical analyses were performed using unpaired, two-tailed Student’s t test (*p= 0.0232, t=3.229, df=5 (E) and ^ns^p= 0.4177, t=0.8828, df=5 (F), ^ns^p= 0.7674, t=0.3123, df=5 (G), ^ns^p= 0.8389, t=0.2128, df=5 (H), ^ns^p= 0.7549, t=0.3298, df=5 (I), ^ns^p= 0.1840, t=1.541, df=5 (J), ^ns^p= 0.5714, t=0.6053, df=5 (K), ^ns^p= 0.5332, t=0.6690, df=5 (L) and ^ns^p= 0.9087, t=0.1206, df=5 (M). Bars represent mean values ±SEM. N=4 and 3 animals for control and *PV-atg5KO-TdT*, respectively.

In contrast, the number of PV-immunoreactive (and Calbindin-negative) interneurons in the cerebellar molecular cell layer was comparable between six-month-old control and *PV-atg5KO* mice (Figure 1C). Similarly, we did not observe any immunoreactivity for active caspase-3 in the cerebellar molecular cell layer of *PV-atg5KO* (Figure S1J, arrowheads, and S1N). Together, these observations invite the speculation that distinct PV-expressing cell populations exhibit differential vulnerability to autophagy perturbation.

In order to assess the requirement of autophagy for PV-neuron maintenance across the brain in an unbiased manner, we then used CLARITY combined with light sheet imaging and machine learning (outlined in Figure 1D) to compare the number of PV-positive neurons between three-month-old *pv-cre;TdT*^*tg/+*^ (control) and *pv-cre;atg5*^*f/f*^*;TdT*^*tg/+*^ *(PV-atg5KO-TdT)* mice. In line with Figure 1A-B, we found a significant reduction in the number of TdT-positive Purkinje cells in the *PV-atg5KO-TdT* brains when compared to controls (Figure 1E), confirming the fidelity of this approach in assessing changes in the number of PV-neurons. Interestingly however, further analysis of eight major brain regions known to contain PV-interneurons indicated no significant differences in the number of TdT-positive cells in *PV-atg5KO* when compared to age-matched controls (Figures 1F-M; expressed as percentage of control in panels 1E-M and as total cell counts in Suppl. Table 1). Furthermore, no significant differences were observed in the volume of these brain structures between the two genotypes (Suppl. Table 1), and a TUNEL assay of three- and six-month-old brain sections confirmed the absence of apoptosis in the *PV-atg5KO* across the forebrain (Suppl. Table 4). Taken together, these results demonstrate that autophagy is not required for the survival of PV-interneurons across the brain.

Given the fundamental role of autophagy in protein turnover, we then speculated that despite not compromising the maintenance of PV interneurons, impaired autophagy may cause deficits in their proteostasis that could directly or indirectly impact their function. To test this hypothesis, we FACS-sorted TdT-positive parvalbumin interneurons from the isocortex and hippocampus of three-month old control (*pv-cre;TdT*^*tg/+*^) and *PV-atg5KO-TdT (pv-cre;atg5*^*f/f*^*;TdT*^*tg/+*^) mice and analyzed them by quantitative proteomics. Principal component analysis indicated a clear segregation of the four control and four knockout replicates based on genotype (Figure S2A). Mass-spectrometry analysis identified a total of 5963 proteins that were detected by at least two unique peptides (Suppl. Table S2). As analysis was performed after FACS-sorting, which can result in breakage of synapses, we first wanted to ensure that synaptic proteins were adequately identified. To this end, we compared our list of identified proteins with the SynGO database and found 824 synaptic proteins present in our list, representing 13.82% of the total proteins detected in our analysis (Figure 2A). The detected synaptic proteins comprised 66.77% of the entire synaptic proteome (Figure S2B), however, did not include any neurotransmitter receptors, presumably due to the limitations posed by FACS-sorting. We also identified 438 mitochondrial proteins, representing 7.35% of the total proteins detected in our analysis (Figure 2A), and 38.42% of the MitoCarta 3.0 mitochondrial protein database (Rath et al., 2021) (Figure S2C). Finally, we detected 392 ER and 534 Golgi proteins, representing 6.6% and 8.96% of the total identified proteins (Figure 2A), respectively, and comprising 74.95% (Figure S2D) and 47.38% of the ER and Golgi proteomes (Figure S2E), respectively (Thul et al., 2017). In order to characterize the effects of autophagy ablation on the PV interneuron proteome, we set a *p*-value of 0.02 (-log10(0.02)=1.7) as the cutoff for determining the proteins that are differentially abundant between control and *PV-atg5KO-TdT* samples. This was based on the enrichment of the positive control p62 (Sqstm1) in the *PV-atg5KO-TdT* sample (Figure 2B). As expected, Atg5 was significantly decreased and Alfy (Wdfy3), another selective aggrephagy receptor, was significantly increased in the *PV-atg5KO-TdT* sample, as expected (Figure 2B). Furthermore, we found 281 proteins that were differentially expressed between the two genotypes, 162 of which were enriched and 119 were less abundant in the *PV-atg5KO-TdT* sample, when compared to the control (Figure 2C). Both the significantly enriched and less abundant proteins included synaptic, mitochondrial, ER and Golgi proteins (as summarized in Figure 2D). Notably, while all the ER proteins found to be dysregulated in autophagy-deficient excitatory neurons (Kuijpers et al., 2021) were detected in our mass-spectrometry analysis (Figure S2F), no significant differences in expression were found between genotypes (Figure S2F,G). Interestingly, this was also the case for three selective ER-phagy receptors, namely Calcoco1, Rtn3 and Atl3. Indeed, only four ER proteins (Avl9, Pggt1b, Prex2, Derl1) were significantly enriched in the *PV-atg5KO-TdT* sample, none of which have known roles in calcium homeostasis. Taken together, these results suggest that contrary to what was found in glutamatergic neurons, ER homeostasis may not be impacted in autophagy-deficient interneurons, as is the case for excitatory glutamatergic neurons.

**Figure 2:**
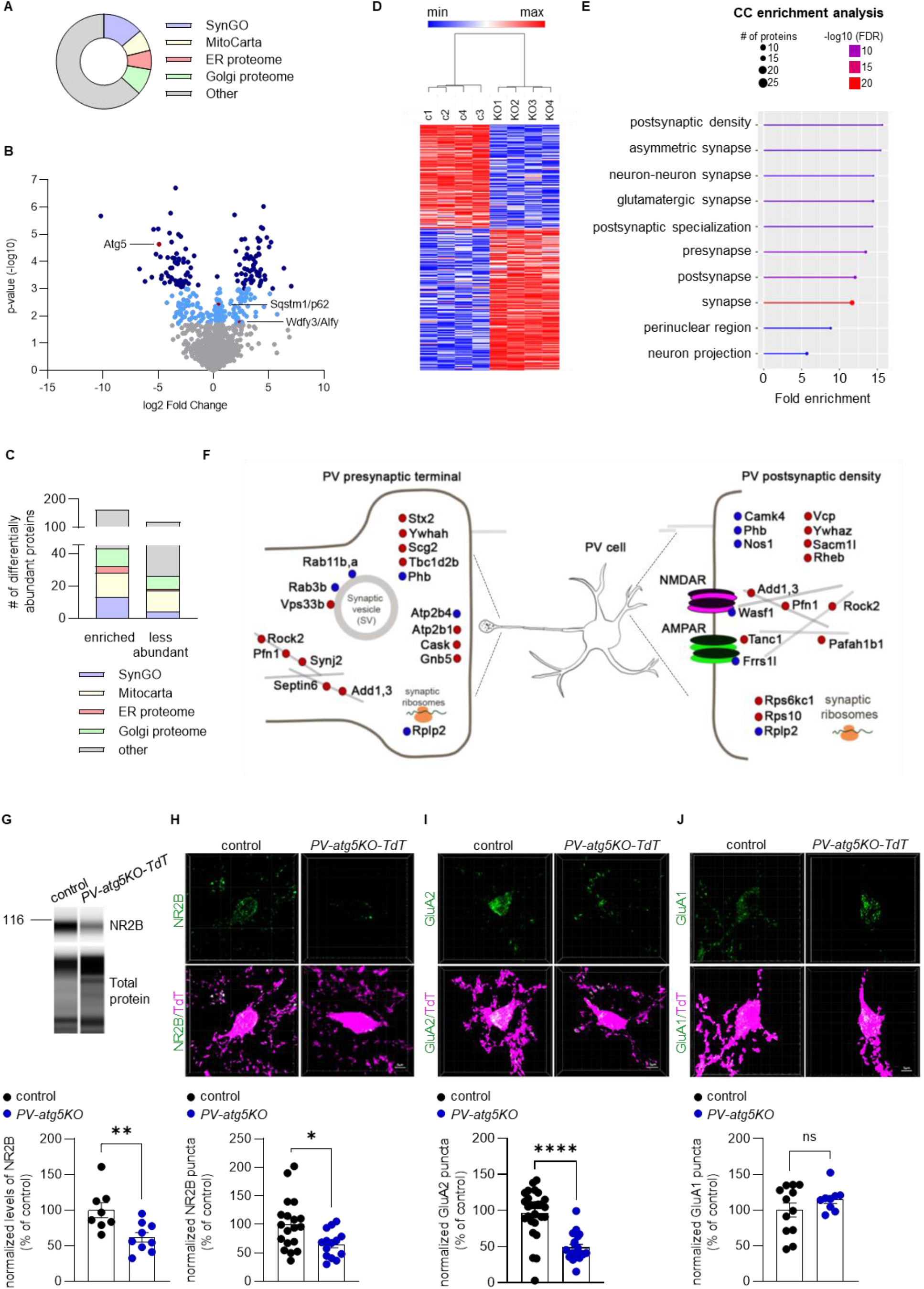
Autophagy regulates the proteostasis of PV-interneurons. A) The part of a whole graph shows the representation of synaptic (13.8%), mitochondrial (7.35%), ER (6.6%) and Golgi (8.9%) proteins in our list of identified proteins, as determined by comparison with the SynGO, MitoCarta, ER and Golgi databases, respectively. B) Volcano plot showing the 5963 proteins which are detectable by proteomics analysis in PV FACS sorted cells (grey dots: -log10(p-value)<1.7, light blue dots: 1.7≤-log10(p-value)<3, dark blue dots: -log10(p-value)≥3; (-log10(0.02)=1.7) and (-log10(0.001)=3). C) Graph showing the differentially expressed proteins including synaptic, mitochondrial, ER and Golgi proteins. Significance was set at p<-log10(p-value)≥1.7; (-log10(0.02)=1.7). D) Heatmap showing the 162 proteins which were found to be enriched and the 119 proteins decreased in the mutant cells, compared to control, showing their hierarchical clustering. Significance was set at p<-log10(p-value)≥1.7; (-log10(0.02)=1.7). E) Graph showing the cellular component enrichment analysis of the deregulated synaptic proteins. F) Graphical representation of the localization of the 32 differentially abundant synaptic proteins in the pre- and post-synaptic compartments of PV-interneurons (red dots: enriched, blue dots: decreased). G) JESS Western blot analysis with an antibody against NR2B in PV FACS sorted cell lysates of three-month-old control *(pv-cre;TdT*^*tg/+*^) and mutant *PV-atg5KO-TdT (pv-cre;atg5*^*f/f*^*;TdT*^*tg/+*^) littermates. Graph showing the protein levels of NR2B normalized to the total protein levels. Statistical analysis was performed using unpaired, two-tailed Student’s t test (**p=0.0072, t=3.109, df=15). Bars represent mean values ±SEM. N=8 and 9 animals for control and *PV-atg5KO-TdT*, respectively. H-J) Immunohistochemistry with antibodies against NR2B (H), GluA2 (I) and GluA1 (J) in CA1 hippocampal area of control *(pv-cre;TdT*^*tg/+*^) and mutant *PV-atg5KO-TdT (pv-cre;atg5*^*f/f*^*;TdT*^*tg/+*^) cells, respectively. PV-interneurons are genetically labeled with Td-tomato (purple). Graphs showing the relative percentage of their surface expression, respectively. Statistical analyses were performed using Student’s t test (*p= 0.0126, t=2.649, df=31, a ****p<0.0001, t=5.69, df=44 and ^ns^p= 0.246, t=1.20, df=19 respectively). Bars represent mean values ±SEM. N=4 animals pergenotype. Scale bar: 5μm.

Comparison of the differentially abundant proteins with the SynGO database indicated that 9 synaptic proteins were significantly downregulated (Wasf1, Atp2b4, Camk4, Frrs1l, Nos1, Phb, Rab11, Rplp2 and Vps33b) and 23 synaptic proteins were significantly enriched (Rps6kc1, Tanc1, Rab3b, Add3, Cask, Tubb2b, Pfn1, Vcp, septin-6, Synj2, Tbc1d2b, Rock2, Ywhah, Gnb5, Atp2b1, Stx2, Scg2, Rheb, Ywhaz, Sacm1l, Pafah1b1, Add1, Rps10) in *PV-atg5KO-TdT* neurons, when compared to control. Cellular component enrichment analysis of the deregulated synaptic proteins indicated that they are mainly localized in the postsynaptic specialization of asymmetric synapses, while some also showed a presynaptic localization (Figure 2E; schematically summarized in Figure 2F). The only deregulated synaptic protein involved in calcium homeostasis was Atp2b4, a plasma membrane calcium ATPase that localizes to the presynaptic membrane, which was found to be significantly decreased in autophagy-deficient PV-interneurons, as confirmed by JESS-Western blot analysis (Figure S2H).

Another deregulated synaptic protein, Wasf1, is a key component of the WAVE protein complex, a key regulator of the actin cytoskeleton formation through interactions with the Arp2/3 complex. In addition, recent work demonstrated that this machinery also links diverse cell surface receptors to the actin cytoskeleton, including the NMDA receptor subunit NR2B (Chen et al., 2014). Thus, we examined whether expression of NR2B is affected in autophagy-deficient PV neurons. To this end, we performed Jess-Western analysis of the TdT-positive PV-interneurons sorted from control or *PV-atg5KO-TdT* mice using an antibody against NR2B and found a significant reduction in its total levels in the autophagy-deficient interneurons when compared to the control (Figure 2G). In addition, we performed NR2B surface immunolabelling on hippocampal sections of control and *PV-atg5KO-TdT* mice and found that autophagy-deficient PV-interneurons exhibit a significant reduction in surface NR2B levels when compared to the control (Figure 2H). In addition, another protein that was significantly downregulated is Frrs1l, an outer component of AMPA receptors (Madeo et al., 2016; Schwenk et al., 2012). With recent work indicating that loss of Frrs1l leads to intracellular retention of GluA2 subunits (Madeo et al., 2016), we also compared the surface expression of GluA2 in hippocampal sections of control and *PV-atg5KO-TdT* mice. We found that autophagy-deficient PV-interneurons exhibited a significant reduction in surface GluA2 levels (Figure 2I). By contrast, surface labeling for the GluA1 subunit showed no differences between genotypes (Figure 2J).

Taken together, these findings indicated that autophagy-deficiency leads to changes in neurotransmitter receptor composition on the membrane of PV-interneurons. We then hypothesized whether these changes in synaptic proteostasis have repercussions on their function, subsequently affecting the networks in which they are embedded. To test this, we focused on the hippocampal CA1 region, where PV expression is restricted to inhibitory interneurons, mostly comprising of basket cells (Donato et al., 2015; Donato et al., 2013). Thus, to examine whether *atg5* deletion influences PV neuronal excitability, we performed patch clamp recordings of current-evoked action potentials from acute brain slices of three-month-old-control (*pv-cre;TdT*^*tg/+*^) and *PV-atg5KO-TdT* (*pv-cre;atg5*^*f/f*^*;TdT*^*tg/+*^) mice (Figure 3A). On average, the input-output curve in PV neurons from slices of *PV-atg5KO-TdT* mice presented a rightward shift compared to their control littermates, indicative of reduced neuronal excitability within this neuronal population (Figure 3A, B).

**Figure 3:**
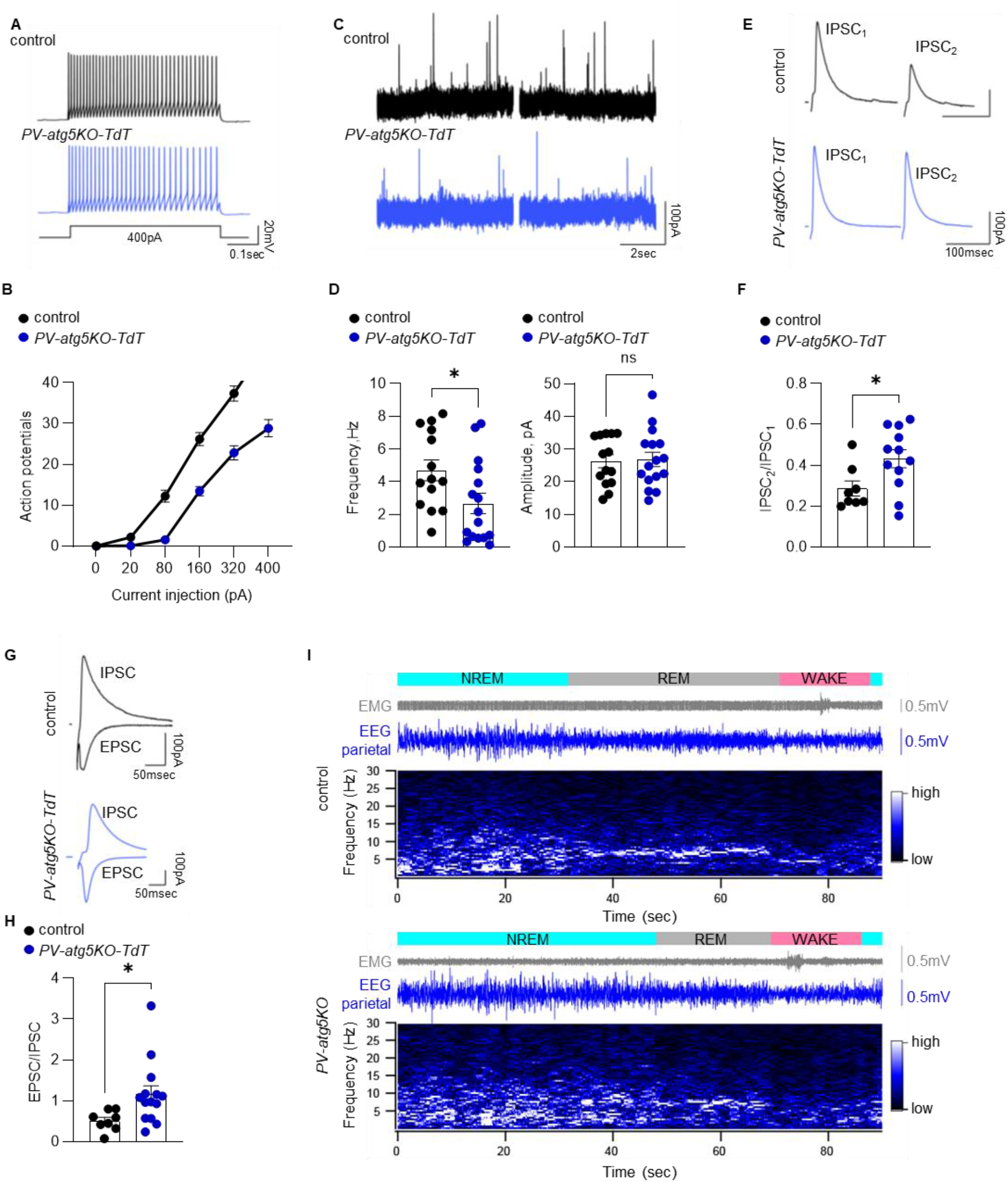
Autophagy-deficient PV-interneurons exhibit functional adaptations leading to E/I imbalance. A) Current-clamp sample traces (400 pA injection, top) and the action potentials evoked by current steps (bottom) from PV neurons, from slices of control *(pv-cre;TdT*^*tg/+*^) and mutant *PV-atg5KO-TdT (pv-cre;atg5*^*f/f*^*;TdT*^*tg/+*^), three-month old mice were recorded. B) Graph showing input-output plot from control *(pv-cre;TdT*^*tg/+*^) (n=14,N=3) and *PV-atg5KO-TdT (pv-cre;atg5*^*f/f*^*;TdT*^*tg/+*^) PV neurons (n=16, N=3). Statistical analysis was performed using two-way repeated measures ANOVA followed by Sidak post hoc test; F(5,140)=23.85 (****p<0.005). Bars represent mean values ±SEM. C) Representative traces of sIPSC from slices of control *(pv-cre;TdT*^*tg/+*^) and *PV-atg5KO-TdT (pv-cre;atg5*^*f/f*^*;TdT*^*tg/+*^) three-month old mice. D) Graphs showing the frequency and amplitude of sIPSCs from control *(pv-cre;TdT*^*tg/+*^) and mutant *PV-atg5KO-TdT (pv-cre;atg5*^*f/f*^*;TdT*^*tg/+*^) (n=21, N=3). Statistical analysis was performed using unpaired, two-tailed Student’s t test (*p=0.0298, t=2.289, df=28 and ^ns^p=0.8654, t=0.1710, df=28, respectively). Bars represent mean values ±SEM. E) Representative traces of paired-pulse ratio (IPSC_2_/IPSC_1_) from three-month old control *(pv-cre;TdT*^*tg/+*^) and *PV-atg5KO-TdT (pv-cre;atg5*^*f/f*^*;TdT*^*tg/+*^) mice. F) Graph showing the paired-pulse ratio (IPSC_2_/IPSC_1_) from control *(pv-cre;TdT*^*rg/+*^) (n=8, N=2) and *PV-atg5KO-TdT (pv-cre;atg5*^*f/f*^*;TdT*^*tg/+*^) (n=12, N=2). Statistical analysis was performed using unpaired, two-tailed Student’s t test (*p=0.0310, t=2.340, df=18). Bars represent mean values ±SEM. G) Representative traces of excitation to inhibition ratio (EPSC/IPSC) from three-month old control *(pv-cre;TdT*^*tg/+*^) and *PV-atg5KO-TdT (pv-cre;atg5*^*f/f*^*;TdT*^*tg/+*^) mice. H) Graph showing the excitation to inhibition ratio (EPSC/IPSC) from three-month old control *(pv-cre;TdT*^*tg/+*^) (n=8, N=2) and *PV-atg5KO-TdT (pv-cre;atg5*^*f/f*^*;TdT*^*tg/+*^) (n=14, N=3) mice. Statistical analysis was performed using unpaired, two-tailed Student’s t test (*p=0.0351, t=2.261, df=20). Bars represent mean values ±SEM. I) Representative 90-sec traces and corresponding time-frequency plots as heatmaps from EEG/EMG recordings from three-month-old control *(atg5*^*f/f*^) and *PV-atg5KO* (*pv-cre;atg5*^*f/f*^) mice.

Next, we examined whether this reduced PV excitability has repercussion on GABAergic neurotransmission onto neighboring pyramidal cells (PC) in the CA1 region. Recording of spontaneous inhibitory postsynaptic currents (sIPSCs) from PC unraveled a reduction in the frequency, but not amplitude of events (Figure 3C, D), concomitantly with an increased paired-pulse ratio of evoked IPSCs (Figure 3E, F). Additionally, we probed excitatory (-60 mV, EPSC) and inhibitory synaptic transmission (+5 mV, IPSC) which revealed enhanced EPSC to IPSC ratio (EPSC/IPSC) in PC from *PV-atg5KO-TdT* mice compared to control littermates (Figure 3G, H). Taken together, these results support a diminished PV neuronal excitability, which accounts for a reduction in synaptic inhibition onto PC.

The enhanced excitation/inhibition ratio onto PC, urged us to further investigate the presence of spontaneous epileptic seizures, as observed in mice with conditional ablation of autophagy in forebrain excitatory neurons (Tang et al., 2014) (Overhoff, 2022) and human patients with mutations in autophagy genes (Fassio et al., 2020). To this end, we performed frontal and parietal electroencephalogram (EEG) recordings in combination with electromyogram (EMG) in three-month-old control (N=8 mice) and *PV-atg5KO* (N=7 mice) animals for three consecutive light-dark phases. Animals cycled between the three vigilance states wake, non-rapid-eye-movement sleep (NREM) and rapid-eye-movement sleep (REM), as evident by standard spectral EEG signatures. Analysis of these recordings using the Epi-AI online peer-reviewed tool (Wei et al., 2021), confirmed the absence of overt spontaneous epileptic seizures in the *PV-atg5KO* mice (Figure 3I), indicating that perturbed autophagy in PV-positive cells may not contribute to epileptic phenotypes.

As PV-interneuron activity has been implicated in memory (Ognjanovski et al., 2017; Tripodi et al., 2018; Xia et al., 2017) and anxiety (Courtin et al., 2014), we then sought to determine whether impaired autophagy affects cognition via changes in PV-interneuron excitability. To this end, three-month-old control and *PV-atg5KO* mice were subjected to a fear conditioning test, a paradigm of associative memory (see Methods, summarized in Figure 4A). While control and *PV-atg5KO* animals exhibited comparable freezing percentages during training, *PV-atg5KO* animals exhibited a significant reduction in freezing on the test day when compared to control littermates (Figure 4B). Differences in freezing cannot be attributed to an impaired ability of *PV-atg5KO* animals to perceive the stimulus since both genotypes showed a significant increase in reactivity upon shock delivery (Figure 4C).

**Figure 4:**
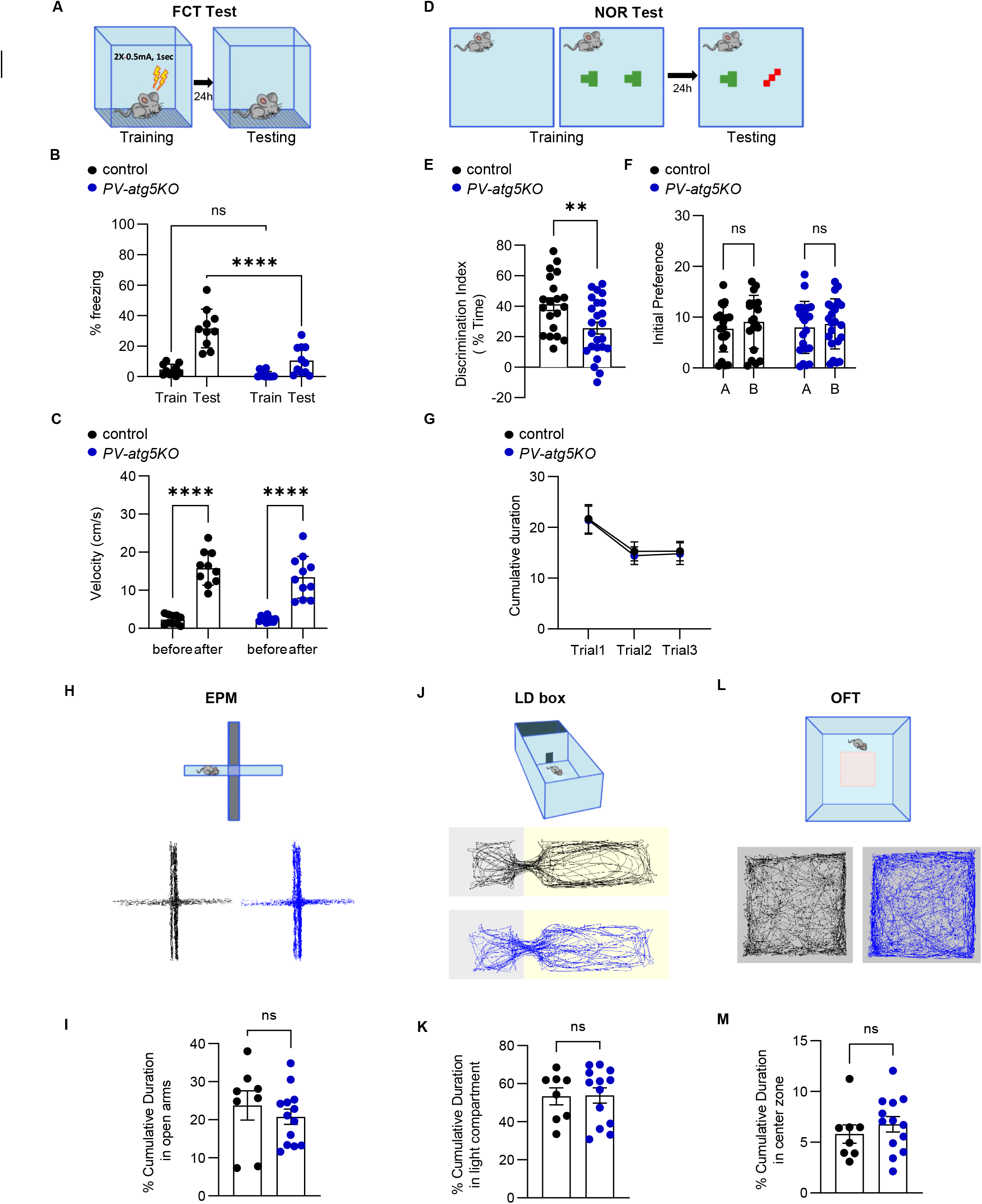
Animals with autophagy deficiency in PV-expressing neurons exhibit deficits in memory. A) Overview of the experimental pipeline for fear conditioning test (FCT). B-C) Graphs showing the percentage of freezing responses and the speed before and after electrical shock in FCT test from three-month old control (*atg5*^*f/f*^) and *PV-atg5KO (pv-cre;atg5*^*f/f*^) male littermates. Statistical analyses were performed using two-way repeated measures ANOVA followed by Sidak post hoc test; F(1,19)=13.17, (train day ^ns^p= 0.5732 and test day ****p <0.0001) and F(1,19)=1.255, (before and after in control ****p <0.0001 and in *PV-atg5KO* ****p <0.0001), respectively. Bars represent mean values ±SEM. N=10 and 11 animals for control and *PV-atg5KO*, respectively. D) Overview of the experimental pipeline for novel object recognition test (NOR). E-G) Graphs showing the discrimination index (DI) in testing day (E) and the initial preference (F) and the cumulative duration (G) spending in objects per trails in training day. Three-month-old control (*atg5*^*f/f*^) and *PV-atg5KO (pv-cre;atg5*^*f/f*^) male littermates were assessed in NOR test. DI was calculating by the following formula: DI= [time in novel object-familiar one/ total time in objects]X100%. Statistical analyses were performed using unpaired, two-tailed Student’s t test (**p=0.0088, t=2.747, df=42) for DI and two-way repeated measures ANOVA followed by Sidak post hoc test for the other two analyses (for initial preference F(1,80)=0.08937, for control ^ns^p=0.6333 and PV-atg5KO ^ns^p=0.8728) and (for cumulative duration F(2,88)=0.02500, for trial 1: ^ns^p=0.9999, for trial 2: ^ns^p= 0.9815 and for trial 3: ^ns^p=0.9974, respectively). Bars represent mean values ±SEM. N=20 and 24 animals for control and *PV-atg5KO*, respectively. H) Overview of the experimental pipeline for elevated plus maze test (EPM) and representative tracking traces from a control (*atg5*^*f/f*^) and a *PV-atg5KO (pv-cre;atg5*^*f/f*^) mouse. I) Graph showing the percentage of cumulative duration in open arms during the EPM test. Two-month-old control (*atg5*^*f/f*^) and mutant *PV-atg5KO (pv-cre;atg5*^*f/f*^) male littermates were assessed in this test. Statistical analysis was performed using unpaired, two-tailed Student’s t test (^ns^p= 0.4565, t=0.7602, df=19). Bars represent mean values ±SEM. N=8 and 13 animals for control and *PV-atg5KO*, respectively. J) Overview of the experimental pipeline for light dark box test (LD box) and representative tracking traces from one control (*atg5*^*f/f*^) and one *PV-atg5KO (pv-cre;atg5*^*f/f*^) mouse. K) Graph showing the percentage of cumulative duration in light compartment during the LD box test. Two-month-old control (*atg5*^*f/f*^) and *PV-atg5KO (pv-cre;atg5*^*f/f*^) male littermates were assessed in this test. Statistical analysis was performed using unpaired, two-tailed Student’s t test (^ns^p= 0.94120.4565, t=0.07473, df=19). Bars represent mean values ±SEM. N=8 and 13 animals for control and *PV-atg5KO*, respectively. L) Overview of the experimental pipeline for open field test (OFT) and representative tracking traces from one control (*atg5*^*f/f*^) and one *PV-atg5KO (pv-cre;atg5*^*f/f*^) mouse. M) Graph showing the percentage of cumulative duration in center zone during the OFT test. Two-month-old control (*atg5*^*f/f*^) and *PV-atg5KO (pv-cre;atg5*^*f/f*^) male littermates were assessed in this test. Statistical analysis was performed using unpaired, two-tailed Student’s t test (^ns^p= 0.4358, t=0.7961, df=19). Bars represent mean values ±SEM. N=8 and 13 animals for control and *PV-atg5KO*, respectively.

To further probe the involvement of autophagy in the regulation of memory processes, control and *PV-atg5KO* mice were subjected to a novel object recognition test, a paradigm of recognition memory (see Methods, summarized in Figure 4D). The difference in time spent with the familiar or the novel object is measured as a discrimination index (DI) and it is a proxy for the memory for the familiar object. *PV-atg5KO* mice exhibited a significant decrease in their discrimination index compared to control littermates, indicating a defective memory for the familiar object (Figure 4E). This deficit was neither due to differences in initial preference (Figure 4F) nor to differences in the cumulative duration spent exploring the objects (Figure 4G). Taken together, these findings indicate that autophagy deficiency in parvalbumin neurons is sufficient to induce profound deficits in both associative and recognition memory.

Lastly, to determine whether autophagy-deficient PV neurons impact anxiety-like behaviors, we tested control and *PV-atg5KO* mice in three anxiety tests: a) the elevated plus maze (EPM), b) the light/dark box (LD), and c) the open field test (OFT) (see Methods, summarized in Figures 4H-L). In all three tests, no significant differences were observed in the cumulative duration that *PV-atg5KO* animals spent in the open arm (Figure 4I), in the light compartment (Figure 4K) or in the center of the arena (Figure 4M) when compared to control littermates. Moreover, the total locomotion of control and *PV-atg5KO* animals was comparable in all three tests (Figure S3A-C). Taken together, these findings suggest that the described proteostatic and synaptic deficits of autophagy-deficient PV neurons, as well as of the networks in which they take part, do not culminate into changes in anxiety-like behaviors.

## Discussion

Emerging evidence from animal models suggests that the well-conserved autophagic machinery, which is fundamental for the turnover of proteins and organelles, is implicated in higher order behaviors that depend on the orchestrated function of multiple neuronal populations. However, our knowledge of the roles of autophagy at cellular and synaptic levels is thus far limited to glutamatergic and other projections neurons. Here we expanded this knowledge to other neuronal populations, demonstrating that autophagy regulates the homeostasis and physiology of parvalbumin-expressing interneurons, safeguarding hippocampal network dynamics and memory.

While the physiological roles of autophagy in interneurons were overlooked, interneurons have already been implicated in the pathology of congenital diseases of autophagy, such as tuberous sclerosis complex (TSC), a disorder that manifests with severe synaptic and psychiatric deficits, including memory deficits (Ridler et al., 2007), epilepsy, developmental delay, and autism (Henske et al., 2016). In this disorder, autosomal dominant germline mutations in *TSC1* or *TSC2* genes lead to the hyperactivation of the mTORC1 pathway, an important positive regulator of protein translation and negative regulator of autophagy. Interestingly, mice with condition ablation of *tsc1* in GABAergic interneuron progenitor cells (Fu et al., 2012) exhibit reduced numbers of GABAergic cells and synapses in the forebrain. Here, we directly tested whether autophagy regulates PV-interneuron numbers and whether it safeguards their maintenance in the brain but found no evidence for such a role. Instead, across all tested brain areas, PV-interneuron numbers where comparable between control mice and those lacking *atg5* conditionally in PV-expressing neurons. Therefore, we conclude that autophagy is dispensable for the survival and maintenance of postmitotic PV-interneurons. We speculate that the interneuron loss observed in the *tsc1*-conditional knockout model may be due to the concomitant deregulation of the translation machinery. Alternatively, it may also be due to the fact that ablation of *tsc1* starts embryonically in GABAergic neuron progenitors, consistent with previous work showing that mTOR signaling regulates the early proliferation of these cells (Ka et al., 2017).

In this study, we directly examined for the first time the requirement of autophagy for inhibitory neurotransmission, focusing on PV-expressing interneurons of the hippocampus. We report that autophagy-deficient PV-interneurons are less excitable, compared to controls. Consistently, we also found reduced inhibitory input to neighboring excitatory neurons of the hippocampus of *PV-atg5KO* animals, compared to control littermates. Together, these findings demonstrate that autophagy-deficient PV-interneurons exhibit reduced basal transmission, which accounts for an imbalance in the excitation-inhibition equilibrium in the hippocampus. Our findings help explain previous work, showing that loss of *tsc1* reduces inhibitory synaptic transmission (Bateup et al., 2013; Khlaifia et al., 2022) and results in fewer and smaller synapses formed by PV-interneurons, which are responsible for social deficits (Amegandjin et al., 2021). As synaptic deficits of PV-interneurons are likely to also pertain to human TSC patients, restoring autophagy specifically in these cells may represent a novel strategy for reinstating inhibitory transmission and network balance, to ameliorate some aspects of the cognitive impairment pathology, such as memory, learning and sociability.

A recent study reported that autophagy-deficient excitatory glutamatergic neurons exhibit enhanced basal transmission, which was attributed to the failed turnover of presynaptic ER and the consequent elevation of intracellular calcium stores (Kuijpers et al., 2021). Taken together with our findings, it emerges that under basal conditions, autophagy regulates excitatory and inhibitory neurotransmission in diametrical ways, restricting the former, whereas enhancing the latter. The mechanisms by which autophagy regulates these two types of transmission may also be distinct. Unlike excitatory neurons, autophagy-deficient PV-interneurons do not exhibit impaired proteostasis of ER proteins or accumulation of selective ER-phagy receptors, although these proteins are well-detected in our proteomic analysis. We cannot exclude that calcium levels are implicated in the reduced inhibitory transmission of autophagy-deficient PV-interneurons, as the proteomic analysis of these neurons revealed significantly decreased levels of Atp2b4, a P-type plasma membrane calcium ATPase (PMCA), which is localized in the presynaptic membrane (Stafford et al., 2017). However, we also found a significant reduction in Frrs1l and Wasf1, which are involved in the plasma membrane retention and targeting of AMPA receptor and NMDA receptor subunits, respectively. This led us to reveal that autophagy indirectly regulates the surface levels of GluA2 and NR2B in PV-interneurons, both of which may be directly implicated in the reduced excitability of these neurons when autophagy is impaired. More generally, it also appears that the autophagic machinery sequesters and turns over distinct cellular substrates in different neuronal populations, adding yet another aspect of selectivity to its contribution to protein and organelle homeostasis.

Finally, we found that while anxiety is unaffected, *PV-atg5KO* animals exhibit deficits in associative and recognition memory, two major components of declarative memory that mainly involve the hippocampus, among other brain areas. Previous work has clearly shown that conditional deletion of AMPA or NMDA receptor subunits in PV neurons causes reduced excitability of fast-spiking PV interneurons in the hippocampus, leading to impaired hippocampal network synchrony and memory deficits (Korotkova et al., 2010) (Fuchs et al., 2007). Consistently, pharmacogenetic inhibition of hippocampal PV-expressing interneurons impairs hippocampal network dynamics, leading to deficits in the consolidation of associative memory (Ognjanovski et al., 2017). Therefore, we believe that the memory deficits of *PV-at*g5KO animals are attributed to the diminished activity of PV-interneurons in the hippocampus and not related to the loss of the cerebellar Purkinje cells, which are instead involved in implicit memories, including procedural memories and motor learning (De Zeeuw and Ten Brinke, 2015).

In conclusion, it emerges that the requirement of autophagy for proper brain function relies on cell-type specific functions in different neuronal subpopulations. Here, we have highlighted the distinct roles played by autophagy in PV-expressing neurons. Whether autophagy regulates the neurotransmission of other types of inhibitory neurons, such as those expressing somatostatin, calretinin, VIP or other markers, remains to be investigated by future studies, in order to gain a complete picture on the collective impact of autophagy impairment on multi-neuronal networks.

## Data availability statement

Requests for resources and reagents should be addressed to the Lead Contact, Vassiliki Nikoletopoulou. All raw MS data together with MaxQuant output tables are available via the Proteomexchange data repository (www.proteomexchange.org). Project Name: Autophagy is dispensable for the maintenance of parvalbumin interneurons but required for their inhibitory neurotransmission and memory. Project accession: PXD035264.

Username

Password: mlWXmW0O

## Author contributions

L.B. performed light sheet microscopy experiments and J.S developed tools for the analysis of these experiments. L.R. designed the behavioral experiments. L.F. and A.L. designed and performed the EEG experiments with help from A.K. M.M. performed all slice electrophysiology experiments. T.C. performed all other experiments and analyses and contributed to writing the manuscript. V.N. led the study and contributed to writing the manuscript. This work was supported by an ERC starting grant (714983-Neurophagy) to V.N., by the SNSF (31003A_175549) to M.M., by the Swiss National Science Foundation (no. 310030-184759 to A.L.) and by Etat de Vaud to A.L.

### Acknowledgements

We thank Maxime Alessandri for his support with the Epi-Ai platform. We also thank Dr. Erin Wosnitzka for critical reading of the manuscript. Behavioral experiments where run in the NeuroBehavioral Analysis Unit (Neuro-BAU) of the Department of Fundamental Neuroscience, FBM/UNIL. We acknowledge all members of the A.L. for help and support.

## Methods

### Animal care

This study was performed in Greece and in Switzerland. Therefore, the animal protocols for the experiments performed in Greece were approved by the FORTH Animal Ethics Committee (FEC) based on the European Union guidelines on animal experimentation, while those performed in Switzerland were approved by the Swiss veterinary services of the Canton Vaud (Licenses VD3460, VD3461). In both cases, special attention was paid to the implementation of the 3Rs, housing, the environmental conditions and analgesia, in order to ensure the animals’ welfare. Mice were housed in Individually Ventilated Cages (IVCs; Innovive, France) in groups of 3-5 per cage. Water and food were provided *ad libitum* with constant temperature (22°C), humidity (55%), air changes (60/hour) and a 12h/12h light/dark cycle (7am ON/7pm OFF), except for mice that are tested in EEG experiment where cycle was 9am ON/9pm OFF). All mice were littermates and were randomly allocated to experimental conditions. A C57BL/6 genetic background of at least for 20 generations was common in all crossings. We worked with the following genetically modified mice: *atg5*^*f/f*^ (Hara et al., 2006), *pv-cre*^*+/-*^ (B6;129P2-Pvalbtm1(cre)Arbr/J-Jax lab # strain 008069) (Hippenmeyer et al., 2007) and a cre reporter TdT^tg/+^ (B6.Cg-Gt(ROSA)26Sortm9(CAG-tdTomato)Hze/J) (Madisen et al., 2010). A*tg5*^*f/f*^ mice were crossed with the *pv-cre*^*+/-*^ mice to ablate *atg5* in parvalbumin positive GABAergic neurons, generating (*pv-cre*^*+/-*^*;atg5*^*f/f*^) *PV-atg5KO mice. pv-cre*^*+/-*^ and *PV-atg5KO* mice were also crossed with TdT^tg/+^ mice to label the PV positive neurons *(pv-cre*^*+/-*^*;atg5*^*f/f*^*;TdT*^*tg/+*^*) PV-atg5KO-TdT*. Male *pv-cre*^*+/-*^ mice were not included in the breeding to exclude any possibility of unwanted germline recombination and global recombination due to the low expression of PV in sperm. The weight of control (*atg5*^*f/f*^), (*pv-cre*^*+/-*^) and *PV-atg5KO (pv-cre*^*+/-*^*;atg5*^*f/f*^) male littermates was monitored systematically and compared at two specific time points, namely at three-weeks and at three-months of age.

### Western Blotting

Western blot was performed as previously described (Nikoletopoulou et al., 2017) with minor modifications. Briefly, tissues were lysed in RIPA Buffer (50mM Tris-Cl pH 7.4, 150mM NaCl, 1mM EDTA, 1% TritonX-100, 0,1% SDS and 0,1% sodium deoxycholate, supplemented with protease inhibitors. Tissues were sonicated for 20 seconds on ice and centrifuged twice for 30min at 16.200g in order to precipitate the DNA. BCA Protein Assay Kit was used to measure the protein concentration following manufacturer’s instructions and the proteins were separated using (7.5-15%) polyacrylamide gel and were transferred to a nitrocellulose membrane which has a 0.2μm pores size (Millipore). The total protein levels were visualized and normalized on the membrane, using the No-Stain(tm) Protein Labeling Reagent, according to the manufacturer’s instruction. Then, the membrane was incubated in 5% skim milk for 1hour and overnight in the primary antibody at 4°C. The membranes were washed three times (3×10min) with TPBS pH 7.4 (137mM NaCl, 2.7mM KCl, 10mM Na_2_HPO_4_, 1.8mM KH_2_PO_4_, 0.1% Tween-20) and were incubated with a secondary horseradish peroxidase-conjugated-HRP antibody (1:10000) for 1h at room temperature. The membranes were washed three times (3×10min) with TPBS and incubated with chemiluminescent solution (SuperSignal West Pico and Femto Substrate-ThermoFisher Scientific) according to the manufacturer’s instruction. The primary antibodies with their dilution which used for western blotting are presented in Table 1. Data were calculated as the optical density ratio (OD) of the sample/total protein. All data were also expressed as the percentages of the average of the control group. Bands were quantified using Fiji Image J software, using the module of analyzing gels.

### Tissue fixation and cryo-protection

For adult brain collection, all adult mice were transcardially perfused with cold 4% PFA (4% PFA in PBS pH 7.4). Then brains were dissected and 3 hours post-fixation at 4°C was performed and then incubated in 30% sucrose in PBS at 4°C until fully penetrated, for cryoprotection. Adult brains were then embedded in Tissue-Tek (Tissue-Tek O.C.T. Compound) on dry ice and stored at 80°C for cryo-sectioning. All brains were cut into coronal or sagittal sections of 16μm and were placed on Superfrost Plus slides. Tissue slices were dried at 37°C for 2-3 hours and stored at -80°C. For immunostaining, the process is described below in methods.

### Immunostaining

Immunostaining was performed as previously described (Nikoletopoulou et al., 2017). In brief, following fixation, 16μm-thick brain sections were rinsed in PBS pH 7.4 (137mM NaCl, 2.7mM KCl, 10mM Na_2_HPO_4_, 1.8mM KH_2_PO_4_) and incubated for 1 hour in blocking solution containing 1% BSA and 0.2-0.5% Triton-X (depending on the antibody), in PBS pH 7.4. Then, the sections were incubated in blocking solution containing primary antibody overnight at 4°C. Blocking solution without primary antibody was applied in separate samples to test for non-specific labeling. The primary antibodies and their final concentrations are summarized in the resource Table 1. Subsequently, sections were rinsed three times (3×10min) in PBS and incubated with the secondary antibodies, prepared in PBS, as indicated in resource table 2, for 1 hour at room temperature. The nuclear dye Hoechst was used (1:5000) to stain nuclei. Tissue was rinsed three times (3×10min) in PBS and mounted onto slides. Confocal images of fluorescently labeled proteins were captured using the LSM 900 (Zeiss). Images were analyzed by Fiji Image J or Imaris Softaware (9.6.1 version).

### TUNEL assay

TUNEL staining was performed using the *in-situ* cell death detection kit (Roche) in sagittal paraffin sections of 4μm. DNase treatment was applied to the tissue as a positive control, following the manufacturer’s instructions.

### Tissue dissociation and FACS

Brains from control *(pv-cre;TdT*^*tg/+*^), mutants *PV-atg5KOtdT (pv-cre*^*-*^*;atg5*^*f/f*^*;TdT*^*tg/+*^) and negative control *(atg5*^*f/f*^*;TdT*^*tg/+*^), three-month-old, were isolated in cold PBS. The isocortex and hippocampal formation were dissected, and individual cells were isolated using the papain dissociation system, following manufacturer’s instruction. Subsequently, myelin elimination was performed using a Percoll gradient. 90% of Percoll in PBS was prepared and the pH was adjusted to 7.4 (Percoll Solution). 3ml of PBS and 1ml of the Percoll Solution was added the myelin-cleared material and mixed well. Then, 4ml of PBS was added to create two phases. Subsequently, the samples were centrifuged at 3000rcf for 10min at 4°C, with acceleration ramp at 1 and braking ramp at 0. The supernatant was discarded and the cell pellet was resuspended in 1ml PBS (Zehnder et al., 2021). The whole process is carried out on ice. Cell sorting (FACS) was performed using a MoFlo AstriosEQ High speed cell sorter.

### CLARITY and 3D image acquisition

Mice were transcardially perfused with 4% PFA and the brain was dissected and was postfixed overnight in 4% PFA. Brains were clarified following the CLARITY protocol (Chung and Deisseroth, 2013), using X-CLARITY™ system. Brains were immersed in a refractive index matching solution (RIMS) containing Histodenz (Sigma Aldrich) for at least 24 hours before being imaged. Imaging was performed using a mesoSPIM system at Wyss Center, Geneva (Voigt et al., 2019). The microscope consists of a dual-sided excitation path using a fiber-coupled multiline laser combiner (405, 488, 561 and 647 nm, Toptica MLE) and a detection path comprising a 42 Olympus MVX-10 zoom macroscope with a 1× objective (Olympus MVPLAPO 1×), a filter wheel (Ludl 96A350), and a scientific CMOS (sCMOS) camera (Hamamatsu Orca Flash 4.0 V3). The excitation paths also contain galvo scanners for light-sheet generation and reduction of shadow artifacts due to absorption of the light-sheet. In addition, the beam waist is scanned using electrically tunable lenses (ETL, Optotune EL-16-40-5D-TC-L) synchronized with the rolling shutter of the sCMOS camera. This axially scanned light-sheet mode (ASLM) leads to a uniform axial resolution across the field-of-view (FOV) of 5μm. Image acquisition is done using custom software written in Python. Z-stacks were acquired with a zoom set at 1.25X at 5μm spacing, respectively resulting in an in-plane pixel size of 5.26 × 5.26μm (2048×2048 pixels). Excitation wavelength of the td-Tomato and autofluorescence channels were set at 561 and 488nm, respectively with an emission filter, 593/40 LP and 530/43 nm bandpass filter respectively (BrightLine HC, AHF). Images were stitched and reconstructed in 3D using Arivis Vision 4D software.

### Registration to the Allen Reference Atlas, Mask Extraction and Segmentation

#### Brain registration to Allen Brain Atlas

Mouse brains were un-anisotropically down-sampled and registered to the 25µm Allen brain mouse atlas (Lein et al., 2007) using the AMAP pipeline (Niedworok et al., 2016) through Brainreg frontend (Tyson et al., 2021) using the autofluorescence channel. The affine registration was computed using 5 steps and the registration was computed for each affine step. The freeform registration was computed using 5 steps and the registration was computed on the last 3 steps only. The bending weight energy was set to 0.9 which was a good tradeoff for accommodating sample imperfections while maintaining high regularization to avoid overfitting the data. All other parameters were kept as default.

#### Mask creation for sub-region segmentation

To accelerate the segmentation in Arivis Vision 4D software, we restricted the volume on which the segmentation was performed by using a binary mask of the volume of interest. To create the binary masks, we used a custom developed tool which interface with the Allen Brain Atlas API and automatically generates any region as a binary mask. This tool allows to create a mask using Allen Brain Atlas region names or acronyms. It automatically and efficiently looks for children’s regions to create masks and allows the user to batch create masks when the input is a list of regions. The output binary masks were produced at 25µm isotropic resolution for memory footprint efficiency and resampled later in Arivis Vision 4D software at the sample resolution for the segmentation step.

#### Segmentation

The masked region was imported into Arivis Vision 4D software and used for the creation of 3D masked region for cell counting analysis. Td-Tomato positive cells (PV-cells) and neuronal fibers were classified using random forest machine learning-based algorithm (ML). Due to the diversity of the brain areas and heterogeneity of background of whole brains scans, the ML algorithm was trained blindly for each brain area and sample. Based on the ML training the 3D objects were segmented and filtered by features, as volume (voxel count) and object sphericity using Arivis Vision 4D software. The raw counted cells were normalized by masked region volume.

### Proteomic Analysis

#### Protein Digestion

Replicate samples were digested according to a modified version of the iST method (Kulak et al., 2014) (named miST method). Briefly, frozen cell pellets were resuspended in 30ul miST lysis buffer (1% Sodium deoxycholate, 100mM Tris pH 8.6, 10mM DTT) by vortexing. Resuspended samples were heated at 95°C for 5min. Samples were then diluted 1:1 (v:v) with water containing 4mM MgCl2 and Benzonase (Merck #70746, 100x dil of stock = 250 Units/μl) and incubated for 15 minutes at RT to digest nucleic acids. Reduced disulfides were alkylated by adding 1/4 vol of 160mM chloroacetamide (final 32mM) and incubating at 25°C for 45min in the dark. Samples were adjusted to 3mM EDTA and digested with 0.2μg Trypsin/LysC mix (Promega #V5073) for 1 hour at 37°C, followed by a second 1 hour digestion with a second and identical aliquot of proteases. To remove sodium deoxycholate, two sample volumes of isopropanol containing 1% TFA were added to the digests, and the samples were desalted on a strong cation exchange (SCX) plate (Oasis MCX; Waters Corp., Milford, MA) by centrifugation. After washing with isopropanol/1% TFA, peptides were eluted in 200ul of 80% MeCN, 19% water, 1% (v/v) ammonia.

#### Liquid Chromatography-tandem Mass spectrometry

Eluates after SCX desalting were dried and resuspended in variable volumes of 0.05% trifluoroacetic acid, 2% acetonitrile to equilibrate concentrations. 1μg of each sample was injected on column for nanoLC-MS analysis.

#### MS analysis

Data-dependent LC-MS/MS analyses of samples were carried out on a Fusion Tribrid Orbitrap mass spectrometer (Thermo Fisher Scientific) interfaced through a nano-electrospray ion source to an Ultimate 3000 RSLCnano HPLC system (Dionex). Peptides were separated on a reversed-phase custom packed 40cm C18 column (75μm ID, 100Å, Reprosil Pur 1.9μm particles, Dr. Maisch, Germany) with a 4-90% acetonitrile gradient in 0.1% formic acid (total time 140min). Full MS survey scans were performed at 120’000 resolution. A data-dependent acquisition method controlled by Xcalibur 4.2 software (Thermo Fisher Scientific) was used that optimized the number of precursors selected (“top speed”) of charge 2+ to 5+ while maintaining a fixed scan cycle of 0.6sec. HCD fragmentation mode was used at a normalized collision energy of 32%, with a precursor isolation window of 1.6m/z, and MS/MS spectra were acquired in the ion trap. Peptides selected for MS/MS were excluded from further fragmentation during 60sec.

#### MS Data analysis

Tandem MS data were processed by the MaxQuant software (version 1.6.14) (Cox and Mann, 2008) incorporating the Andromeda search engine (Cox et al., 2011). The UniProt Mus musculus reference proteome (RefProt) database of August 26th, 2020 was used (55’508 sequences), supplemented with sequences of common contaminants. Trypsin (cleavage at K,R) was used as the enzyme definition, allowing 2 missed cleavages. Carbamidomethylation of cysteine was specified as a fixed modification. N-terminal acetylation of protein and oxidation of methionine were specified as variable modifications. All identifications were filtered at 1% FDR at both the peptide and protein levels with default MaxQuant parameters. MaxQuant data were further processed with Perseus software (Tyanova et al., 2016). iBAQ (Schwanhausser et al., 2011) values were used for quality control assessment, after which LFQ values (Cox et al., 2011) were used for quantitation after log2 transformation. Only protein groups were retained which had all four valid values in at least one condition. In the resulting table, missing values were imputed with values determined from a normal distribution (parameters used in Perseus: width 0.3 and down-shift 2.2 SD).

### Jess-WB analysis

Cell pellets were resuspended in SDS lysis buffer (50mM Tris pH 6.8, 2% SDS, 5% glycerol, 2mM DTT, 2.5mM EDTA, 2.5mM EGTA, 4mM NaVO4, 20mM NaF complemented with Complete Protease inhibitors and PhosStop (Roche)) and heated 10min at 65°C. Then, 4.8μl of protein extracts were subjected to the capillary-based immunoassay Jess system (ProteinSimple) using the 12-230kDa separation module (ProteinSimple SM-W004) with the Replex (RP-001) and the Total Protein (DM-TP01) modules according to the manufacturer’s instructions. Analyses were performed using the Compass software version 6.1. Peaks were determined using the dropped line method and data were normalized to the Total Protein.

### In vitro electrophysiology

The mice 11-13 weeks were anesthetized (ketamine/xylazine; 150mg/100mg/kg), sacrificed, and their brains were transferred in ice-cold carbogenated (95% O_2_/5% CO_2_) solution, containing (in mM) choline chloride 110; glucose 25; NaHCO_3_ 25; MgCl_2_ 7; ascorbic acid 11.6; sodium pyruvate 3.1; KCl 2.5; NaH_2_PO_4_ 1.25; CaCl 20.5. Coronal brain slices (250μm thickness) were prepared and transferred for 5min to warmed solution (34°C) of identical composition, before transfer at room temperature in carbogenated artificial cerebrospinal fluid (ACSF) containing (in mM) NaCl 124; NaHCO_3_ 26.2; glucose 11; KCl 2.5; CaCl_2_ 2.5; MgCl_2_ 1.3; NaH_2_PO_4_ 1. During recordings, slices were immersed in ACSF and continuously super fused at a flow rate of 2.5mL min-1 at 32°C. Neurons were patch-clamped using borosilicate glass pipettes (2.7–4MΩ; Phymep, France) under an Olympus-BX51 microscope (Olympus, France). Signal was amplified, filtered at 5kHz and digitized at 10kHz (Multiclamp 200B; Molecular Devices, USA). Data were acquired using Igor Pro with NIDAQ tools (Wavemetrics, USA). Access resistance was continuously monitored with a -4mV step delivered at 0.1Hz. Extracellular stimulation from AMPI ISO-Flex stimulator was delivered through glass electrodes placed nearby the pyramidal cell patched. Current-clamp recordings were obtained using an internal solution containing (in mM): potassium gluconate 140; KCl 5; HEPES 10; EGTA 0.2; MgCl_2_ 2; Na_2_ATP 4; Na_3_GTP 0.3 mM; creatine phosphate 10. Current-clamp experiments were performed by a series of current steps (negative to positive) injected to induce action potentials (-20, 0, 20, 80, 160, 320, 400pA injection current, 800ms). For recordings in voltage-clamp configuration, evoked EPSCs were recorded at -60mV while evoked and spontaneous inhibitory currents were recorded at +5mV. Internal solution contained (in mM) CsMeSO_3_ 120, CsCl 10, HEPES 10, EGTA 10, creatine phosphate 5; Na_2_ATP 4; Na_3_GTP 0.4, QX-314 5. All drugs were purchased from HelloBio.

### Polysomnographic EEG/EMG recordings

Control (*atg5*^*f/f*^) and *PV-atg5KO (pv-cre;atg5*^*f/f*^) male littermates, three-month-old, were used under a 12h/12h light/dark cycle (light onset at 9:00 AM). EEG and EMG electrodes were implanted as described (Fernandez et al., 2017). Briefly, animals underwent a gas anesthesia (1-2 % isoflurane with O_2_) and gold-plated electrodes were implanted over *the dura mater* through frontal and parietal bones in order to acquire frontal and parietal EEG recordings. Two EMG electrodes (gold pellets) were inserted into the neck muscles. A silver wire (Harvard Apparatus) implanted in the bone of cerebellum was used as ground and neutral reference to record referential EEG signals. Electrodes were glued to the skull (cyanoacrylate), soldered to a male-to-male connector, and covered with dental cement (Palavit, Heraeus Kulzer GmbH).

Paracetamol (2mg/ml) was diluted into the drinking water for 7 days of recovery after the surgery, and an additional week of adaptation to the freely moving cabling system. Animals were monitored according to a score sheet established with the Veterinarian Authorities. Signals were recorded in freely moving conditions with animals in their home cage, amplified, digitized and acquired (RHD2132 amplifier chip, connected to a RHD USB interface board C3100 from Intan Technologies, Los Angeles, CA) at 1000Hz sampling frequency via Matlab recording software and Intan Matlab toolbox.

### Seizure and epileptic events detection

For the seizure and epileptic event detection, we used the following free online software (https://lifescience.ucd.ie/Epi-AI/), as described (Wei et al., 2021).

### Behavioral analyses

#### Open field test (OFT)

Two-month-old control (*atg5*^*f/f*^) and *PV-atg5KO (pv-cre*^*-*^*;atg5*^*f/f*^) male littermates were habituated in the behavioral room in single cages for at least 1 hour before each experiment. Anxiety-like behavior was assessed in a PVC square open field arena (L=45cm, W=45cm and H=30cm) with gray opaque walls, where a central square zone was virtually marked 15cm from the four walls of the arena. The ambient illumination was 20lx in the chamber (measurement without infrared light). Infrared light (850nm) was used during the experiment to improve the contrast and tracking detection. Mice were placed for 20min in the arena and their body-center was tracked with EthoVision XT15 (Noldus Information Technology, Wageningen, The Netherlands). Total distance (cm), velocity (cm/s), number of entries in the center zone, and the percentage of time spent in the center zone were scored. Mice were tested between 10:00am and 3:00pm.

#### Light/Dark test

Two-month-old control (*atg5*^*f/f*^) and *PV-atg5KO (pv-cre*^*-*^*;atg5*^*f/f*^) male littermates were tested for their preference for the dark or light compartment of a two-chambered arena, using the Noldus Light/Dark set up. In this set up, the dark compartment (L=20cm, W=20cm, H=20 cm) is smaller than the light compartment (L=40cm, W=20cm, H=20cm). The two compartments were separated by a manually-operated sliding door. The walls and ceiling of the dark compartment were permeable to infrared light, allowing optimal mouse tracking. The ambient illumination was 3-4lx in the dark and 600-700lx in the light compartment (measurement without infrared light). Mice were habituated in single cages inside the experimental room for at least 1 hour prior the test.

During the test, every mouse was placed in the dark compartment with the door closed. After 10sec, the sliding door was opened manually, allowing free transition between compartments for 5min. The frequency to shuttle between compartments and the time spent in each compartment were scored using EthoVision XT15 (Noldus Information Technologies). The frequency of transition between the areas, the duration, the total distance (cm) and the velocity (cm/s) were measured throughout the test session. Mice were tested between 10:00am and 3:00pm.

#### Elevated Plus Maze (EPM)

Two-month-old control (*atg5*^*f/f*^) and *PV-atg5KO (pv-cre*^*-*^*;atg5*^*f/f*^) male littermates, were subjected to Elevated Plus Maze (EPM) for evaluation of anxiety-like behavior. The EPM (Noldus Information Technologies) is a cross-shaped arena elevated from the floor (H:47cm) consisting of two open and two closed arms (dimensions of each arm: L=36cm, W=6cm). The arms are infra-red backlit to optimize the tracking of the mouse in closed arms. The ambient illumination was measured in the closed arm (4-6lx), in the center (40-60lx), and in the open arm (104-108lx). The mice were habituated in single cages inside the testing room for 1 hour before the test. Each mouse was individually placed in the center of the EPM facing one of the close arms and was allowed to freely explore the maze for 5min. The number of transitions between the open and closed arms, the percent time spent inside the zones of the maze, the total distance traveled (cm), and the velocity (cm/s) were monitored with EthoVision XT15 (Noldus Information Technologies). Anxiety-like was measured as an increase in closed arms activity (either duration or number entries). Mice were tested between 10:00am and 3:00pm.

#### Spontaneous activity Test

Spontaneous activity of two to three-month-old Control (*atg5*^*f/f*^) and *PV-atg5KO (pv-cre;atg5*^*f/f*^) male littermates was measured in the PhenoTyper 3000 (Noldus Information Technologies). The PhenoTyper 3000 is a home-cage for mice (L:30cm x W:30cm) optimized for video tracking, via EthoVisionXT14 (Noldus Information Technologies). Mice were placed in the Phenotypers during the light phase (2:00-3:00pm) and were housed in this cage for 72 hours without any further human handling. Regular food and water were provided *ad libitum* during this time. Video tracking started at 7:00pm. A set of spontaneous behavior parameters (described in (Loos et al., 2014) were collected: Food and water consumption, locomotion, sheltering, dark light index, dark light phase transition.

#### Rotarod Test

Three-month-old control (*atg5*^*f/f*^) and *PV-atg5KO (pv-cre;atg5*^*f/f*^) male littermates were subjected to the rotarod test in order to evaluate their motor coordination and balance as previously described (Dominguez-Iturza et al., 2019). Each mouse was habituated in a single cage inside the experimental room for at least 1 hour. The ambient illumination of the experimental room was 25-30lx. Mice were habituated on the rotarod apparatus for 2min at constant speed (4rpm). After 10min, they were tested on rotarod for 3 trials of 5min with an inter-trial interval of 10 min. The speed of rotarod was accelerated from 4rpm to 40rpm during the 5-min trial. The latency to fall off from the rod was recorded. The average latency to fall as well as the latency to fall per trial were measured in these 5min trials. Mice were tested between 10.00am and 3:00pm.

#### Novel Object Recognition Test (NOR)

Three-month-old control (*atg5*^*f/f*^) and *PV-atg5KO (pv-cre;atg5*^*f/f*^) male littermates were subjected to a two-day NOR test. On each day, mice were habituated in single cages inside the experimental room, at least 1 hour before testing. The first day, mice are individually placed inside an open arena (L=45cm, W=45 cm and H=30cm) with gray PVC opaque walls in dim light conditions (20lx). The arena was filled with bedding (2cm) in all trials. On the first day, the mouse was habituated inside the empty arena for two 10-min-trials. During the following three trials (inter trial interval= 5min), two identical objects (lego) were placed in the arena and the mouse was free to explore the objects for 10min. At the end of third object exploration trial, the mouse was brought back to its own home cage in the animal vivarium. Twenty-four hours later, mice were individually placed inside the arena for one trial with one familiar object and a novel object. The objects were located at the same distance from each wall (15cm). Mouse activity was tracked with EthoVision XT 14 software (Noldus Information Technologies). The distance travelled per trial, the time spent exploring each object, the object exploration time per each trial, and the discrimination index were measured. Discrimination index was calculated using the following formula: Discrimination Index (%) = [time spent exploring the novel object new - time spent exploring the familiar one/ total time spending in objects]X100%. Mice were tested between 10:00am and 3:00pm.

#### Fear Conditioning Test

Three-month-old control (*atg5*^*f/f*^) and *PV-atg5KO (pv-cre;atg5*^*f/f*^) male littermates were subjected to a two-day Contextual Fear Conditioning test. Mice were first habituated in single cages inside the experimental room at least 1 hour before testing. Each mouse was placed individually in the fear conditioning chamber (Ugo Basile, srl), which is composed of 4 transparent Plexiglas walls and an electrified metal grid floor. After 2 minutes of free exploration, two mild foot-shocks (0.5mA, 2sec) separated by one minute interval, were delivered through the grid-floor. Thirty seconds after the termination of the second foot-shock, the mouse was placed back to its own home cage. Memory for the association between the context and the aversive stimulus was probed 24 hours later for 5min inside the same conditioning chamber used during the training session. An overhead camera connected to the software Ethovision XT15 (Noldus Information Technology) automatically monitors freezing behavior (i.e. crouched posture and absence of any movement except respiratory movements) throughout the training and test sessions. Percentage of time spent freezing during the test session is used as a proxy measure for the memory of the association formed during the training session. The current of the foot shock was measured in every experiment to ensure that it corresponds to 0.5mA. Mice were tested between 10:00am and 3:00pm.

### Statistical analysis

Analyses were performed via GraphPad Prism software for behavioral test and for molecular and biochemical analysis. The statistical tests are reported in the respective figure legend. Briefly, unpaired two-tailed Student’s t-tests were used for comparison between two groups. For comparison of two groups and two variances, two-way ANOVA, Multiple-t-test with Tukey or Bonferroni or Holm-Sidak’s multiple comparison correction was used among groups. For all data analysis, p<0.05 was considered as significant and reported as follows: *p<0.05, **p≤0.01, ***p≤0.001 and ****p≤0.0001. Results were presented as mean and standard error of mean (±SEM).

### Power Analysis

An a priori power analysis was conducted using G*Power version 3.1 (Faul et al., 2007) to determine the sample size required to test differences between control and *PV-atg5KO* mice in anxiety-like tests (EPM, LD box and OFT). Results showed the minimum sample size to achieve 80% power 1-β=0.8 (probability of Type II error) for detecting a medium effect, at a significance criterion of α=0.05 (probability of Type I error) and analyzed with a two-tailed Student’s t test. The results are presented in the Suppl. Table 5.

### Resources and reagents table

**Table 1:** Used chemicals and kit were listed in this table

| **Chemical** | **Supplier** | **Cat #** |
| --- | --- | --- |
| cOmplete™, Mini, EDTA-free Protease Inhibitor Cocktail | MERCK | 11836170001 |
| BCA Protein Assay Kit | Life Technologies Europe BV | 23225 |
| No Stain protein labeling reagent | Life Technologies Europe BV | A44449 |
| Hoechst 33342 | HelloBio | HB0787 |
| Tissue-Tek O.C.T. Compound | Sysmex | 4583 |
| FD Rapid GolgiStain™ Kit | Holzel Diagnostika | PK401-C |
| In Situ Cell Death Detection Kit, Fluorescein | Roche | 11684795910 |
| DNase I Solution | STEMCELL | 7900 |
| Papain Dissociation System | BioConcept (Worthington Biochemical Corporation) | LK003150 |
| Histodenz™ | MERCK | D2158 |
| X-CLARITY™ Hydrogel Solution | KitLogos Byosystem | C1310X |
| SDS Sodium Dodecyl Sulfate | Brunschwig | 45900-0010 |

### Antibody table

**Table 2:** Used antibodies were listed in this table.

**Antibody table***Table 2: Used antibodies were listed in this table.*
| **Antibody** | **Company, Cat #** | **Dilution (process)** |
| --- | --- | --- |
| LC3B-rabbit | Sigma,L7543 | 1:1000 (WB, IHC) |
| LC3B-mouse | Santa cruz-sc,271625 | 1:1000 (WB, IHC) |
| p6 SQSTM1 (p62)-mouse | Calbiochem, DR1057 | 1:5000 (WB) |
| SQSTM1 (p62)- g.pig | Progen, GP62-C | 1:2000 (IHC) |
| ALFY (WDFY3, B-4) | Santa cruz-sc, sc-514569 | 1:1000 (WB) |
| Gephyrin-mouse | Synaptic Systems, SYSY147021 | 1:500 (WB, IHC) |
| Parvalbumin-mouse | Swant, 235 | 1:1000 (WB, IHC) |
| Parvalbumin-rabbit | Swant, PV27 | 1:2000 (WB, IHC) |
| Calbindin D-28k-rabbit | Swant, CB-38a | 1:5000 (IHC) |
| Active-Caspase 3-rabbit | R&D systems, AF835 | 1:200 (IHC) |
| RFP-rabbit | Rockland, 42896 | 1:1000 (WB, IHC) |
| GABA<sub>A</sub> receptor γ2 subunit-rabbit | Synaptic Systems, 224003 | 1:1000 (WB, IHC) |
| vGAT-g.pig | Synaptic Systems, 131004 | 1:500 (WB, IHC) |
| GAD2/GAD65-mouse | Synaptic Systems, 198111 | 1:500 (WB, IHC) |
| NMDAR2A (GluN2A)-rabbit | Alomone (AGC-002) | 1:500 (WB, IHC) |
| NMDAR2B (GluN2B)-rabbit | Alomone (AGC-003) | 1:500 (WB, IHC) |
| GluA2 C-terminus-mouse | Synaptic Systems, 182211 | 1:100 (WB) |
| GluA2 N-terminus-gpig | Alomone (AGP-073) | 1:100 (IHC) |
| ATP2B4-rabbit | Sigma (SAB2100184) | 1:500 (WB) |
| b3-Tubulin/Tuj1-mouse | Santa cruz-sc, 80005 | 1:5000 (WB, IHC) |
| b3-Tubulin/Tuj1-rabbit | Abcam, ab18207 | 1:1000 (WB, IHC) |
| Alexa 488-rabbit | ab150077 | 1:1000 (IHC) |
| Alexa 594-mouse | ab150116 | 1:1000 (IHC) |
| Alexa 647-g.pig | ab150187 | 1:1000 (IHC) |
| IgG (Dylight 800 4xPEG)-mouse | Invitrogen, SA5-35521 | 1:10000 (WB) |
| IgG (H+L)-rabbit | Jackson, 211-035-109 | 1:10000 (WB) |
| IgG (H+L)-mouse | Jackson, 715-035-150 | 1:10000 (WB) |
| Rabbit HRP | Biotechne, DM-001 | 1X, (Jess WB) |
| Mouse HRP | Biotechne, DM-002 | 1X, (Jess WB) |

---

**Supplementary Table 5:**
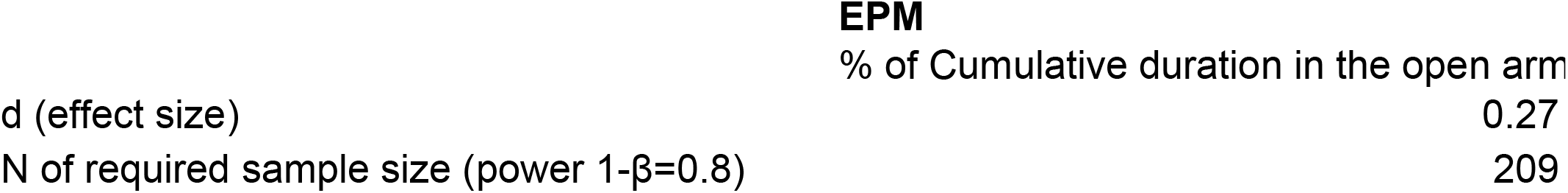
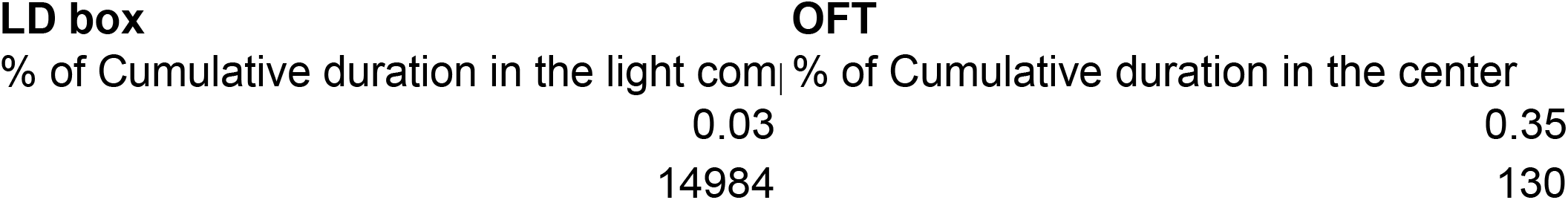
Power Analysis.

